# The Tonoplast Topology Index - a new metric for describing vacuole organization

**DOI:** 10.1101/2025.08.06.668875

**Authors:** Helena Kočová, George Alexandru Caldarescu, Radek Bezvoda, Fatima Cvrčková

## Abstract

**Background:** The plant vacuole arises by orchestrated interplay of membrane trafficking, cytoskeletal rearrangements and a variety of signalling pathways. In the root, the characteristic large central vacuole develops by endomembrane reorganization occurring mainly in the transition zone. The vacuole’s bounding membrane - the tonoplast - can be visualized *in vivo* using fluorescent protein markers, allowing for quantitative analysis of confocal microscopy images. Tonoplast organization can thus serve as a sensitive indicator of changes to any of the processes involved in vacuole biogenesis. The Vacuolar Morphology Index (VMI) is widely accepted as a quantitative measure of vacuole structure. However, this metric has two drawbacks - it only reflects the size of the largest vacuolar compartment (missing therefore possible differences in the organization of smaller compartments), and its determination is labor intensive, limiting its use on large datasets.

**Results:** We developed an alternative metric for describing vacuole organization, named the Tonoplast Topology Index (TTI), which overcomes the above-mentioned shortcomings of the VMI. We compared the performance of our protocol with VMI on a simulated dataset and on real data. To validate the methods’ performance, we used it to confirm the previously reported differences in vacuole shape and size between Arabidopsis *thaliana* roots grown on the surface of an agar medium compared to those embedded inside the agar. Both VMI and TTI could efficiently detect the relatively subtle changes in vacuole organization depending on the position of the root in the agar, and provided correlated results. However, only TTI produced data with close to normal value distribution, simplifying subsequent statistical evaluation.

**Conclusions:** We present the protocol for TTI determination as a two-stage semi-automated procedure involving microscopic image analysis employing an ImageJ macro and subsequent processing of numeric data in the Jupyter Notebook environment, together with benchmarking image data. Since this implementation is freeware-based, platform-independent and (relatively) user-friendly, we hope it will find its use as a high throughput, added value alternative to the VMI metric.

## Background

Plant vacuoles are the largest intracellular membrane compartments, participating in various cellular processes including storage or degradation of cellular components, ion homeostasis or cell expansion (see, e.g., [1]). The vacuoles can attain various shapes, ranging from a highly fragmented collection of smaller compartments or a tubular network in meristematic cells [2,3] up to one big central vacuole with a few invaginations or occasional cytoplasmic strands, which can take up to 90% of the cell volume in more mature expanding or differentiating cells (reviewed in [4]).

In typical angiosperm roots (such as those of *Arabidopsis thaliana*), establishment of the large central vacuole takes place in the transition zone and is a prerequisite for rapid turgor-driven cell expansion. The central vacuole arises by fusion and/or expansion of precursor compartment in a process that involves trafficking of membranes originating from the endoplasmic reticulum, depends on the integrity of the actin cytoskeleton and is intimately linked to auxin signalling (e.g. [2,3,5-7]. Any changes to this process may be thus a source of variability in tonoplast organization in otherwise similarly-sized cells.

Vacuole shape may be affected by various conditions, such as cell type [8], pH [9], sucrose availability and concentration in the media [10], as well as by stiffness of root surroundings, with differences reported between roots penetrating through media compared to those growing on the agar surface [9]. Various mutations or pharmacological treatments can modulate vacuole organization as well (summarized in [11]) although in most cases plant cells somehow manage to complete central vacuole development. However, mutations disabling the molecular machinery critically important for vacuolar development, for example subunits of the HOPS complex [12], are lethal at the embryo stage.

Since the tonoplast can be easily visualized by confocal microscopy using fluorescent dyes or tonoplast-resident proteins fused to various fluorescent proteins (e.g. [3,13,14], quantitative description of tonoplast morphology in differentiating root tissues, whose cells exhibit a relatively regular shape, may serve as a good indicator for following the function of cellular mechanisms and pathways engaged in vacuole biogenesis. A “vacuolar morphology index” (VMI), defined as the product of the length and width of the largest vacuole compartment section in an optical section of a cell (**Fig. 1a**), has been introduced as a quantitative metric for describing the state of vacuole development [6]. This metric has been widely adopted in subsequent studies (e.g., [7,9,15-18].

**Figure 1.**
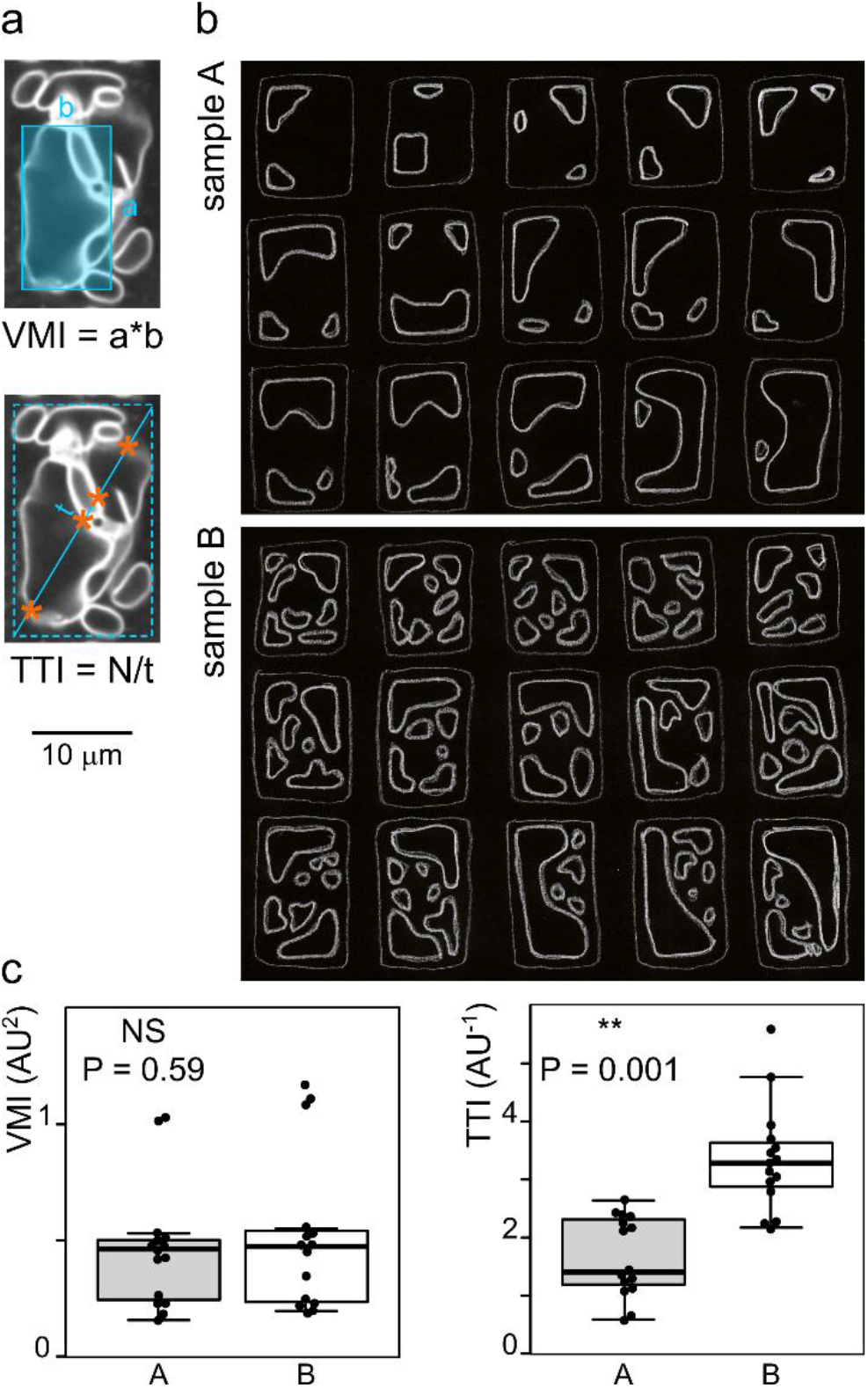
Comparison of the vacuolar morphology index (VMI) and tonoplast topology index (TTI) metrics. (a) A representative optical section of an Arabidopsis transition/elongation zone rhizodermis cell expressing GFP-tagged vacuolar H+ pyrophosphatase with schematic drawing of the principle of VMI and TTI determination. N is the number of membrane crossings along a transect (marked by asterisks). Dashed rectangle is an approximation of the cell outline. (b) Hand-drawn simulated images of two populations of cell optical sections with similar VMI but different vacuole organization. (c) Result of manual VMI and TTI determination from simulated image data shown in (b), documenting that only TTI can distinguish between the two populations. (t-test P values shown; AU - arbitrary units).

However, while the VMI captures well the overall progress of central vacuole enlargement, it cannot, in principle, provide any information about the organization of vacuolar compartments outside the largest (nascent) central vacuole. In this study, we are proposing an alternative metric of vacuole organization, which overcomes this drawback of the VMI, specifically focusing on the overall extent of vacuole fragmentation and complexity of vacuole shape. This method has been tested on simulated data and on a benchmark set of images taken from Arabidopsis roots expressing GFP-tagged vacuolar H+ pyrophosphatase [14] that were grown on the surface or inside agar medium, reproducing a previously reported observation [9]. Together with comparison of TTI and VMI results on these datasets, we are presenting a user-friendly, platform-independent, freeware-based protocol for semi-automated, relatively high throughput TTI determination.

## Methods

### Plants and culture conditions

An *Arabidopsis thaliana* line expressing plant vacuolar H+ pyrophosphatase 1 fused with monomeric GFP (VHP1:mGFP), obtained by crossing an original Col-0 background line [14] to the Col-8 background, was used in all experiments.

Seedlings were grown *in vitro* on vertically positioned Petri dishes containing ½ Murashige and Skoog (MS) medium with 1.6% (w/v) agar and 1% (w/v) sucrose, pH adjusted to 5.7, at 22 °C with 16 h light/8 h dark cycle as described previously [19] for 5 days prior to imaging. Seedlings grown on the media surface were directly mounted into a microscopy chamber, while seedlings with roots growing inside agar media were carefully pulled out from agar before mounting.

### Imaging

Confocal Z-stack images were acquired using a vertical spinning disc confocal microscopy system consisting of a Zeiss Axio Observer 7 microscope integrated with a Yokogawa CSU-W1-T2 spinning disk unit featuring 50 um pinholes and a VSHOM1000 excitation homogenizer (Visitron Systems), along with VisiView software (Visitron System, v4.40.14) and alpha Plan-Apochromat 100×/1.46 Oil immersion objective. A 488 nm laser was employed for GFP (excitation 488 nm, emission 500-550 nm).

### Image processing and image analysis

Routine image processing, VMI determination and the first stage of the TTI determination procedure (including its development) has been performed on a personal computer running the Fiji distribution [20] of ImageJ 1.54p under a 64bit version of Windows 10 or Windows 11 (Enterprise/Pro or Home edition).

Prior to analysis, raw confocal image stacks produced by the microscope software have been processed in Fiji first by algorithmic contrast enhancement to ensure good visibility of the tonoplast. For the purpose of long-time archiving, the image stacks were subsequently converted to 8 bit, rotated to position the root vertically, cropped to decrease resulting file size, and converted to the *.ome.tiff format. All image quantification procedures have been successfully performed both on raw or contrast-enhanced images or on files that underwent processing for archivation, with practically identical results obtained for a random image sample. Quantitative analyses were in all cases performed on atrichoblasts of the rhizodermis, whose vacuole organization differs from that of the trichoblasts [21].

VMI was determined by multiplying manually measured values of the length and width of the largest vacuole compartment found across the optical sections, preferably in (but not limited to) the perinuclear section, i.e. above the nucleus and below the very cortex [6], obtained using built-in Fiji functions. For any given root, all atrichoblasts with suitably positioned optical sections were analyzed (in some cases using more than one optical section); repeated measurements of the same cell on two sections or in two microscope fields were avoided.

### Data processing and statistics

For numeric data processing in the course of TTI protocol development and TTI determination, a Windows PC running Jupyter Notebook as a part of the Anaconda 2.5.3 package has been used together with scripts written in the Python language version 3.11.7. Raw TTI data processing was performed using Scipy library [22] v. 1.11.4. The Matplotlib library [23] v. 3.8.0 was used to generate graphs, while data tables were produced using the pandas library v. 2.1.4 [24,25]. Simple data processing was performed in MS Excel.

R version 4.4.2 [26] was used for plotting, statistical analysis, and additional data processing. The ggplot2 library was utilized for plotting, while data processing was conducted using dplyr library, both part of the Tidyverse package [27]. Some plots were produced also using the R-based BoxPlotR tool [28] or Matplotlib 3.6.3 [23] employing Seaborn 0.13.2 [29]. The Shapiro-Wilk test (α = 0.05; [30]) was applied to assess data distribution. Data that did not conform to the normal distribution were analyzed using the Mann-Whitney test [31], while Student’s t test [32] was used for normally distributed data.

## Results

### Rationale for developing the TTI metric

The VMI metric relies solely on measuring the size of the largest vacuolar compartment crossed by the optical section under study (**Fig. 1a**; see Methods for details of optical section selection). It is, however, conceivable that cells with large vacuoles of similar size and shape may differ, possibly dramatically, in the presence, number and organization of additional smaller vacuolar compartments, as illustrated by a hand-drawn simulated dataset comprising two populations with similar large vacuoles but dissimilar organization of smaller ones (**Fig. 1b**). In spite of a readily visible difference between these populations, there is no significant difference between their average VMI values (**Fig. 1c**).

We propose a new metric, the Tonoplast Topology Index (TTI), that should be able to capture not only the progress of the large vacuole development reflected in the VMI, but also the overall complexity of vacuole shape. We are reasoning that this parameter should be reflected in the number of tonoplast intersections with a defined linear transect, for example one corresponding to the cell diagonal, and that the number of these crossings normalized to the transect length to compensate for cell size variation can serve as a metric of vacuole shape complexity. We thus defined the TTI as the number of tonoplast intersections per unit of diagonal transect length (**Fig. 1a**). Indeed, TTI determination on the simulated dataset using visual peak counting and transect length measurements in Fiji revealed a significant difference between the two model populations (**Fig. 1c**).

### A semi-automated pipeline for TTI determination

While the TTI can be determined manually, partial automation of the procedure can substantially speed up the process while enhancing reproducibility and decreasing the probability of human-introduced errors. We thus developed a two-stage protocol for determining the TTI (**Fig. 2a**), where the first stage involves manual transect definition with simultaneous cell length measurement, and subsequently employs Fiji’s built-in Plot profile function to measure and export intensity profiles of cell transects. The following second stage, performed in the Jupyter notebook environment, analyses the exported profile data to identify peaks corresponding to membrane crossings, counts the peaks, divides the peak count by transect length and outputs a table containing matched cell length and TTI values for individual cells. We are providing the necessary scripts and detailed, beginner-friendly instructions for performing the procedure as **Additional File 1**.

**Figure 2.**
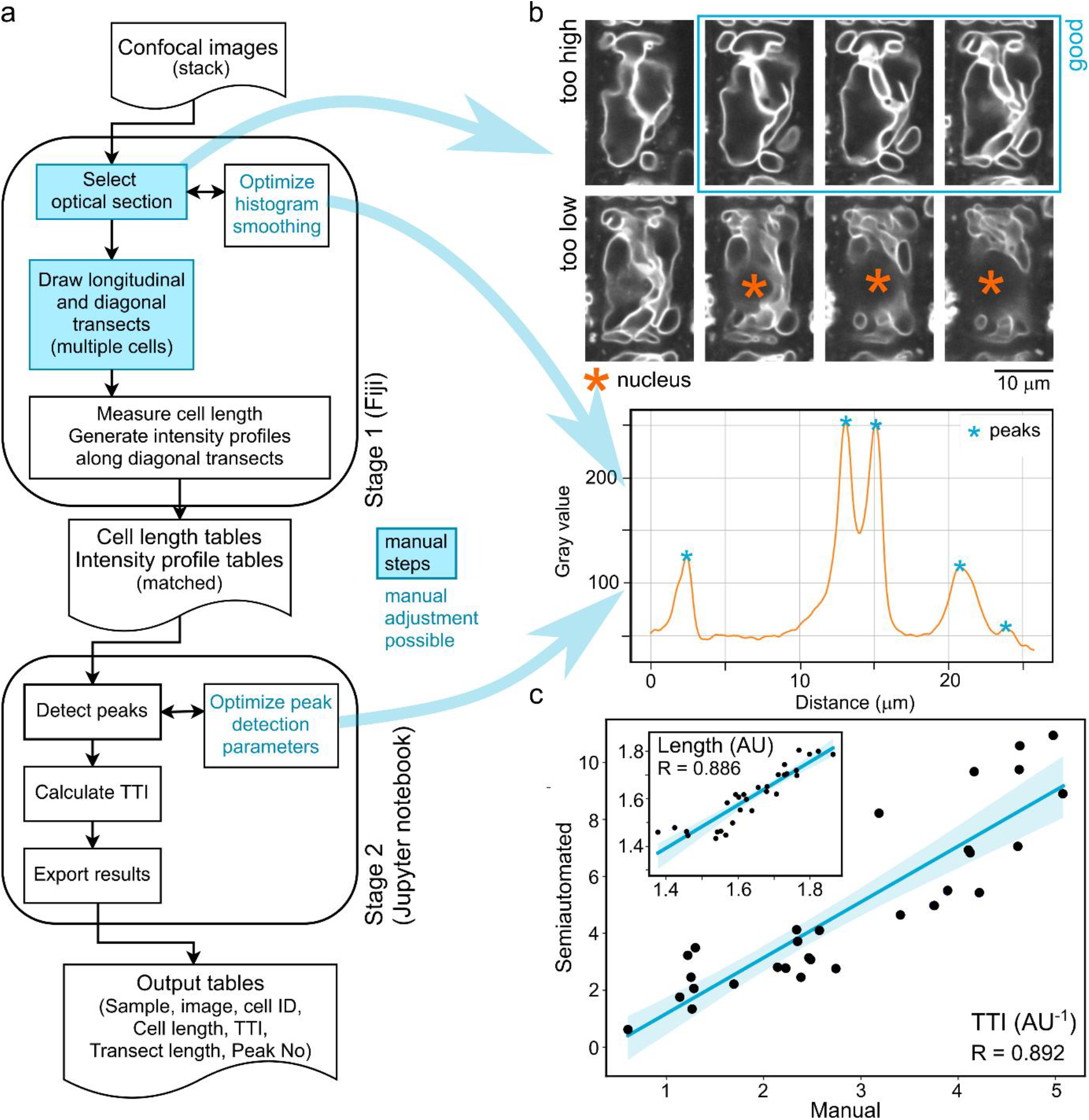
Semi-automated TTI determination procedure. (a) Schematic of the TTI determination pipeline. (b) Examples of optical section selection (top, suitable sections highlighted; the middle one of these is at the optimal position), and acceptable histogram smoothing and peak detection (bottom). Both examples correspond to the cell and diagonal transect shown in Fig. 1a. (c) Comparison of results of manual and semi-automated TTI determination on the simulated dataset from Fig. 1b (with results of cell size measurements shown as an inset). The same transect direction was used for both approaches.

Both stages involve steps that either require manual intervention or include an opportunity to perform manual parameter adjustments. At the first (Fiji) stage, first of them is the selection of an appropriate image from the confocal stack. Measurement of comparable focal planes is crucial, as the tonoplast varies greatly across the cell’s Z dimension. Similar to VMI determination, the measurements should be performed in the perinuclear section, a few micrometers above the nucleus, which appears as a roundish area devoid of tonoplast membranes. Typically, suitable optical sections are located within the top half of the cell’s Z dimension but below its top third (**Fig. 2b**).

Second manual step is the selection of cells to be analyzed and positioning of longitudinal and diagonal transects of each cell for determining cell length and TTI, respectively. The diagonal transects should be oriented uniformly to avoid observer-induced bias, although this may not be critically important for large datasets (see below). Care has to be taken to avoid repeated measurements of the same cell at several planes of focus or image fields.

Third, the intensity profile generation step involves image denoising by repeated rounds of smoothing to improve the signal to noise ratio for subsequent peak detection while preserving information about membrane topology. The number of smoothing rounds can be adjusted depending on the quality of microscopy images, with the preset value optimized for our benchmark data set (see below) but with a possibility of modification (**Fig. 2b**; see also detailed manual in **Additional File 1**).

Lastly, the second (Jupyter notebook) stage of the procedure involves application of several filters to prevent detection of most false positive peaks (local maxima close to the background) while preserving major peaks corresponding to genuine membrane crossings (**Fig. 2b**). The default values of several parameters (**Table 1**) are optimized for our benchmark data but adjustment may be required for a different microscope, tonoplast labeling technique, or image noise level. It is advisable to optimize the stage 1 denoising, as well as the stage 2 filtering, on a subset of source images prior to analysing large datasets.

**Table 1.** Overview of adjustable parameters of Stage 2.

| Parameter | Description |
| --- | --- |
| background | Filters out all peaks below the set value. Useful in case of a very noisy background. If increasing this value, all analyzed plots should have similar intensity ranges to avoid loss of peaks in low intensity plots. |
| med_ratio | Multiplier of the median intensity values from each separate profile, used to filter out maxima lower than median*med_ratio value. Useful to compensate for varying intensity ranges among different profiles, not affected by changing bit depth of the source images. Typically around 1 or slightly lower. |
| min_ratio | Multiplier of the minimum intensity values from each separate profile, used to filter out maxima lower than minimum*min_ratio value. Useful to compensate for varying intensity among different profiles, not affected by changing bit depth of the source images. Typically higher than 1. |
| distance | Minimum distance between two maxima to be considered as separate membrane crossings (scale-independent, takes next/previous value in plot as 1). Value can be adjusted to avoid artificial peak clusters coming from a single membrane. |
| prominence | Minimum height of a peak compared to the closest local minima. Adjust to avoid loss of minor peaks next to major peaks. |
| sigma | Gaussian smooth. Can be used for further denoising in addition to stage 1 smoothing to eliminate artifactual minor peaks. |

To test the performance of our semi-automated pipeline, we used it to analyze our simulated dataset (**Fig, 1b**) and examined the correlation between results of manual and semi-automated TTI determination for individual cells. Although we noticed that the semi-automated method tends to overestimate the TTI due to detecting weak satellite lines on many of the hand-drawn vacuole contours (which is unlikely to occur on real-life microscopy data), we found an overall strong correlation (close to very strong *sensu* [33] between results of manual and semi-automated analysis for both the TTI and cell length (**Fig. 2c**).

### A benchmark data set for TTI procedure validation

In order to validate the TTI determination procedure, we decided to test it by reproducing the previously described difference in vacuolar organization in atrichoblasts of roots grown on agar media surface compared to those penetrating the agar media, with cells from surface-grown roots having generally higher VMI than those from roots embedded in the medium [9]. We were able to obtain high quality confocal images from both types of roots, further referred to as the “top” (surface-grown) and “deep” (agar-embedded) population. Following the published report [9], we focused only on the atrichoblasts, which were readily distinguishable from trichoblasts in the late meristematic zone by their less fragmented vacuolar compartments and overall weaker tonoplast signal (**Fig. 3a**).

**Figure 3.**
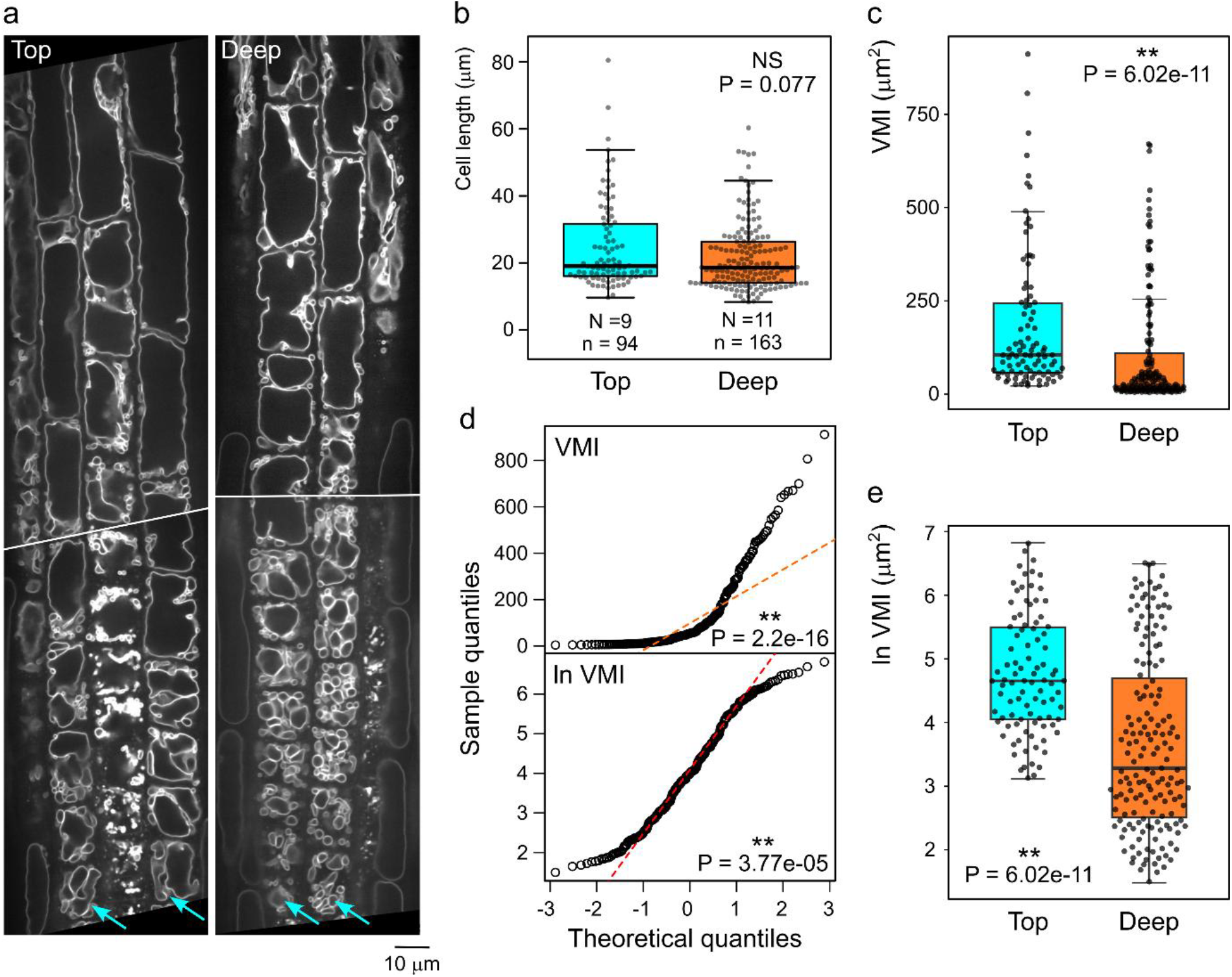
Characterization of the benchmark data set using the VMI metrics. (a) Examples of optical sections of transition and elongation zone rhizodermis of a root grown on the agar surface (labelled “top”) or embedded inside the agar medium (labelled “deep”). Rows of atrichoblasts are marked by arrows. (b) Atrichoblast length distribution for a set of transition and elongation zone cells from top and deep agar-grown roots (N denotes the number of roots, n the number of cells). (c) Comparison of atrichoblast VMI distribution for the cell set from (b). (d) QQ plots of pooled VMI values from top and deep agar showing that the log-transformedVMI value distribution for the dataset from (c) deviates from normal distribution less than that of raw data. Shapiro-Wilk P-values are shown. (e) Comparison of log-transformed atrichoblast VMI distribution for the same data as in (c). Mann-Whitney P-values are shown in (b), (c) and (e).

The atrichoblasts of both populations included in our analyses exhibited a similar asymmetric cell length distribution; although the average cell length was somewhat smaller in the “deep” group, the difference was not statistically significant (**Fig. 3b**). The distribution of VMI values of both populations was even more noticeably asymmetric (**Fig. 3c**), and significantly deviating from the normal distribution. A quantile-quantile (QQ) plot suggested a possible exponential distribution. Indeed, logarithmic transformation brought the VMI value distribution somewhat closer to normal, with its shape possibly corresponding to the Cauchy distribution (**Fig. 3d**). Therefore, the Mann-Whitney test had to be used to assess the statistical significance of observed VMI differences. For both original and log-transformed data (**Fig. 3c,e**), we found significantly higher VMI values in the “top” population compared to the “deep” one.

We thus successfully confirmed that our image data set behaves in agreement with the previous report that cells from roots penetrating into agar media possess more fragmented vacuoles compared to those from surface-grown ones [9], and proceeded to use the same images (also deposited in the EMBL-EBI BioImage Archive repository, accession number S-BIAD2226) as a benchmark set to test the performance of our TTI procedure.

### Comparison of the TTI and VMI metrics on the benchmark data set

We have first used the semi-automated TTI procedure on a subset of 70 cells from the “top” population of the benchmark data set to examine the effect of the diagonal transect direction on the analysis outcome. There was no difference between average TTI values due to transect direction (**Fig. 4a**), indicating that there is no effect of cell chirality and that the transect direction does not affect the analysis outcome, although it should still be kept consistent within any given analysis to avoid observer-introduced bias.

**Figure 4.**
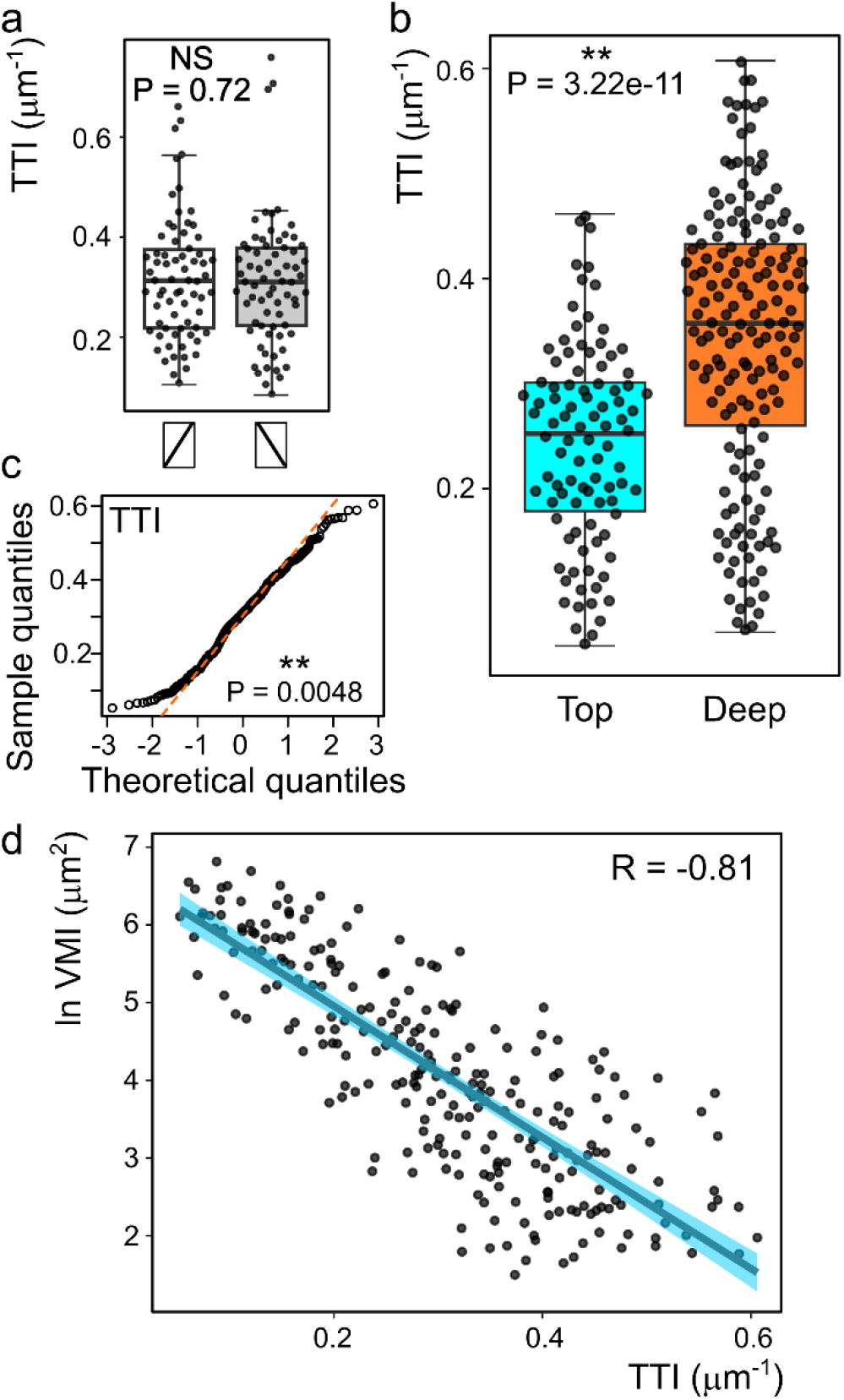
Testing the TTI method on the benchmark data set. (a) Comparison of TTI values obtained on a subset of the benchmark data set using the indicated directions of the diagonal transect (Mann-Whitney P value shown). (b) Comparison of atrichoblast TTI distribution among roots grown on top of the agar medium or deep inside (data from the same set of cells as in Figure 3 b-e, Mann-Whitney P value shown). (c) QQ plots of pooled TTI values from top and deep agar indicating near-normal distribution. Shapiro-Wilk P-value shown. (d) Pearson correlation of TTI and log-transformed VMI values for individual cells from the benchmark data set, indicating negative correlation between the two metrics.

Analysis of the complete benchmark image set using the TTI pipeline revealed significantly higher TTI values (and thus more complex tonoplast topology, as already suggested by VMI determination results) in cells from agar-embedded roots compared to those grown on the media surface (**Fig. 4b**). TTI value distribution was near-normal but not normal, possibly corresponding to the Couchy distribution (**Fig. 4c**), justifying the use of the Mann-Whitney test for statistical evaluation.

Both methods thus documented higher vacuole shape complexity, associated with smaller size of individual vacuolar compartments, in the “deep” population compared to the “top” one. Furthermore, comparison of TTI and log-transformed VMI values for the same cells revealed a strong negative correlation between the results of both methods (**Fig. 4d**), showing that they produce mutually consistent results.

To further examine the effects of root growth conditions on vacuolar organization by both methods, we divided the benchmark cell population into subgroups according to cell length. Splitting the population into quartiles based on cell length (with quartile border values determined from pooled “top” and “deep” populations) allowed us to compare cells not only of similar size, but also of approximately similar developmental age (**Fig. 5**). Independent from the measurement method, significant differences were confined to the first three quartiles (Q1-Q3), corresponding to smaller cells located in the late meristematic and transition zones, while large (Q4) cells did not exhibit statistically significant differences. Thus, the effects of root growth conditions are only detected in small cells but diminish or disappear later due to developmental progression and the expansion of the central vacuole. Importantly, both the conventional VMI method and the newly proposed TTI metric are comparably sensitive with respect to detection of vacuolar organization differences present in our benchmark image data set.

**Figure 5.**
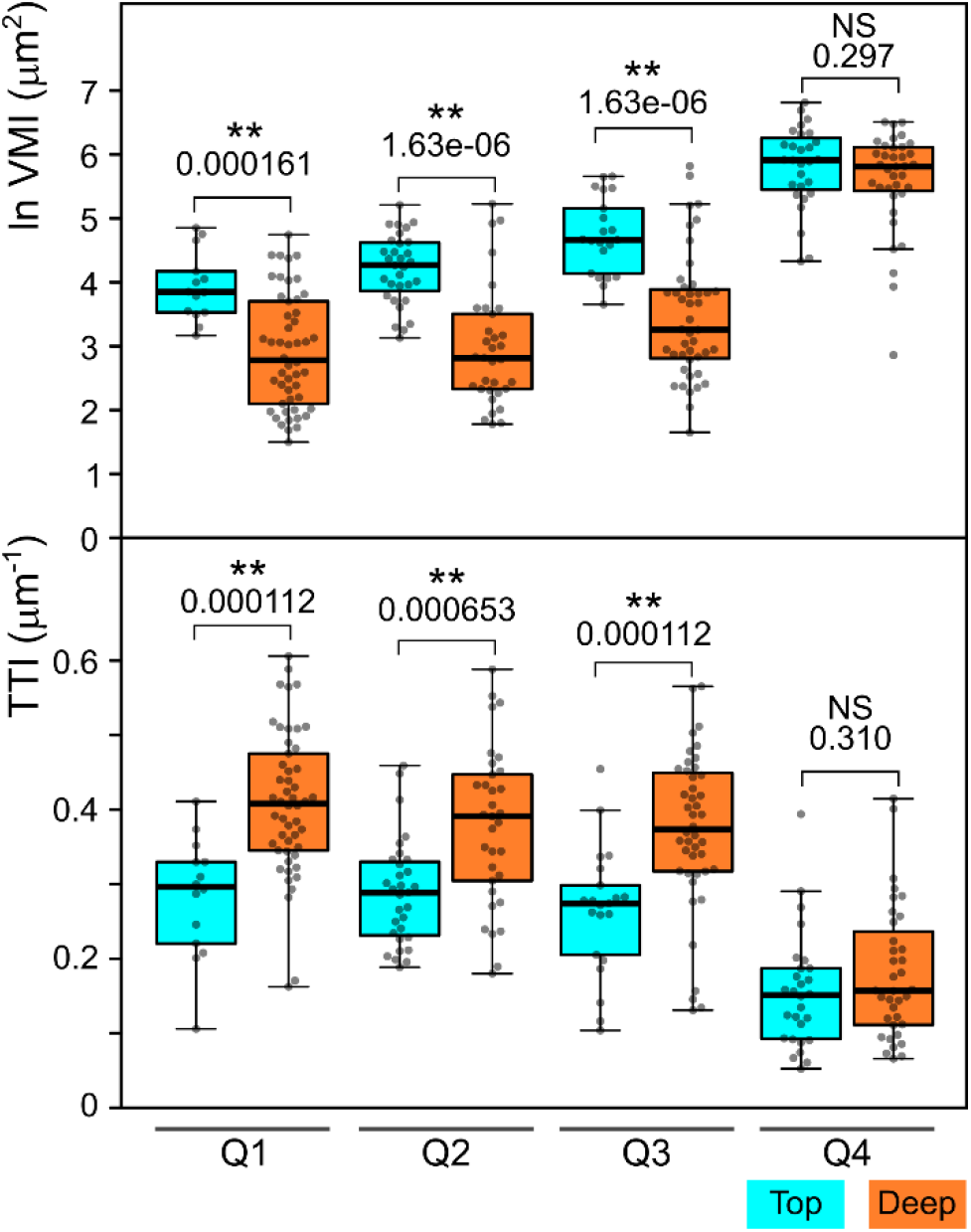
Comparison of TTI and VMI metrics for size-matched cells. Top: Log- transformed VMI value distribution in atrichoblast populations from top and deep agar-grown roots (the same dataset as in Figures 3 and 4); Bottom: TTI value distribution for the same set of cells. For both plots, the cell population was split into subsets corresponding to quartiles of pooled cell length distribution. Mann-Whitney P-values, separately calculated for the indicated pairwise distribution and subjected to Benjamni-Hochberg correction for multiplicity, are shown.

## Discussion

Studies investigating vacuole organization often rely on qualitative descriptions or 3D modelling, typically restricted to a specific cell types such as stomata guard cells [34], tobacco BY-2 cells [35], or rhizodermal cells of distinct zones of the root tip (see Introduction; reviewed also in [11]). While qualitative description may be sufficient in cases where visualisation of the structures itself brings novel insights, i.e., where morphological differences are clearly distinguishable visually, quantitative approaches are necessary to address more subtle phenotypes, especially in cells with complex vacuolar structure, such as those found in root tissues.

Although in recent years reports using quantitative metrics of tonoplast morphology evaluation began to emerge (see Introduction and also, e.g., [36]), the use of quantitative approaches is complicated by technical complexity and computational demands of high data volume image analysis. The Vacuolar Morphology Index (VMI, [6]) has recently gained popularity as an accessible option especially for following vacuole development in root tissues, as it does not require proprietary software or advanced computational skills while being able to detect morphological differences across conditions, although it may not capture all aspects of tonoplast organization, as it focuses solely on the size of the largest vacuole compartment detected in a given cell.

Several alternatives to VMI have been introduced but did not achieve such widespread acceptance. Examples include vacuole volume, surface to volume ratio, or vacuolar occupancy, i.e. the vacuole to cell volume ratio [7], determined in the open-source MorphoGraphX 3D segmentation and quantification environment [37] or in the commercial Imaris software [38,39]. Despite increasing availability of relevant software in microscopy facilities, transferring these tools (and relevant skills) to diverse experimental environments may be quite challenging. A “vacuole shape index” defined as the ratio between the width and length of the largest vacuole compartment (and therefore sharing the detection limits of the VMI) has been recently used alongside VMI (for unclear reasons renamed to “vacuole area”), and occupancy [10], and vacuole compactness has been quantified by measuring the tonoplast to plasmalemma distance at cell corners [15]. Overall, there is so far an obvious lack of standardized workflows for quantitative description of vacuole organization, and development of a new, simple metric that can be scaled up for large datasets may be beneficial.

Here we present the Tonoplast Topology Index (TTI), a new metric that provides a complementary quantitative perspective on vacuolar architecture that is capable of detecting additional aspects of vacuole organization, since it also reflects the complexity of smaller sub-vacuolar compartments. Our protocol is amenable to upscaling, somewhat reduces the requirement for subjective observer’s decisions (such as the choice of the largest vacuolar compartment) that could be introducing artifacts, and may represent an alternative or valuable addition to currently available options.

While establishing a benchmark data set for testing our method, we successfully reproduced the previously reported difference in vacuolar organization in the rhizodermis of roots grown on the surface of agar media compared to those embedded inside [9]. However, characterization of our benchmark data set using the conventional VMI metric alerted us to the notably asymmetric distribution of VMI values in real-life rhizodermis cell populations that is, in our view, only rarely reflected in reported vacuole organization analyses. Multiple studies [6,7,15,17,40] report statistical analyses performed using Student’s t-test on VMI data that are unlikely to meet assumptions of normality and equal variance, and in some cases even display clear asymmetry or variance differences that are not accounted for [15,17]. Non-parametric statistical methods that do not assume normal data distribution, although clearly more appropriate, were used only in some cases [9,16,41]. Vacuolar morphology differences reported in some of the previous studies should thus be interpreted with caution. Although the importance of choosing a non-parametric test for data deviating from normal distribution may be reduced for very large samples, typically consisting of hundreds of individual values [42,43], most studies employ a relatively small number of individual plants per sample and cells per plant; we could find only a single report involving 100 or more cells per sample [15].

In contrast to VMI, our new TTI method generates data exhibiting a distribution that is generally symmetric and relatively close to normal. At the same time, the semi-automated procedure enables large scale analyses, and collection of several hundred data points per sample, though laborious, is possible (if enough good quality confocal images can be collected). Thus, while use of non-parametric methods for statistical evaluation is still recommended for non-normally distributed data, with a sufficiently large sample (see [43]), parametric tests could be employed. For an example, ANOVA can be used if multiple treatments, genotypes or combinations thereof are compared, substantially simplifying data analysis.

Thus, our new TTI metric can contribute towards a more transparent and reproducible framework for quantitative analysis of tonoplast organization. Moreover, while originally intended for measurements of vacuole shape complexity, the method might be suitable for adaptation for other applications involving quantification of geometrically complex biological structures.

## Ethics approval and consent to participate

Not applicable (this manuscript contains no research involving the use of animal or human data or tissue).

## Consent for publication

Not applicable.

## Availability of data and materials

Image data generated and analysed in the current study are available in the EMBL-EBI BioImage Archive repository, accession number S-BIAD2226 (doi: 10.6019/S-BIAD2226, https://www.ebi.ac.uk/biostudies/bioimages/studies/S-BIAD2226). Additional data generated or analysed during this study, including the first release of the software tool developed, are included in this published article and its supplementary information. Archive copy and possible future updates of the software tool generated here are also available at https://github.com/GeorgeCaldarescu/TTI-Tonoplast-Topology-Index.

## Competing interests

The authors declare that they have no competing interests.

## Funding

This work has been supported by the Czech Science Foundation grant 22-33471S (FC, HK, RB) and by the project TowArds Next GENeration Crops, reg. no. CZ.02.01.01/00/22_008/0004581 of the ERDF Programme Johannes Amos Comenius (GAC; part of FC and HK salary during final stages of manuscript preparation).

## Authors’ contributions

H.K and F.C. conceived the project., H.K. obtained the experimental data, G.A.C. and H.K. developed the software, R.B. managed, processed and curated data, H.K., G.A.C. and F.C.analyzed data, F.C. and H.K. drafted the manuscript and figures, G.A.C. and R.B.contributed parts of the text or figures, F.C. performed final editing.

## Acknowledgements

We thank Shoji Segami for sharing the VHP1:mGFP plant line, Falco Kruger and Melanie Krebs (University of Heidelberg) for providing its seeds, Marta čadyová and Dagmar Tůmová for technical assistance, and Matyáš Fendrych for sharing imaging equipment.

## Additional information

**Additional file 1.** Archive file in *.zip format containing scripts and a detailed manual for performing TTI determination.

## Notes

https://github.com/GeorgeCaldarescu/TTI-Tonoplast-Topology-Index

https://www.ebi.ac.uk/biostudies/bioimages/studies/S-BIAD2226

### Competing Interest Statement

The authors have declared no competing interest.

