## Additional file 1 for "The Tonoplast Topology Index - a new metric for describing vacuole organization": stage1_v1.ijm

```
// images are open??

if (nImages == 0){
	exit("No open image. Please open an image to proceed.")
}

// get the path and name of the opened image
inputPath = getDirectory("image");
imageName = getTitle();
baseName = substring(imageName, 0, lastIndexOf(imageName, "."));
// create results folder
outputPath = inputPath + "results/";
File.makeDirectory(outputPath);
run("Duplicate...", "use");

// Retrieve list of existing files
fileList = getFileList(outputPath);

zCount = 1; // start with z1 


// ------ Preprocessing ----------
// dialog box to ask how many times to run Smooth
numSmooth = getNumber("Enter the number of times to apply Smooth:", 2);


// apply smooth the specified number of times
for (i = 0; i < numSmooth; i++) {
	run("Smooth");
}


// show the plot profile
roiManager("select", 0);
run("Plot Profile");

// ------ End of Preprocessing ----------

// ------ Transition Dialog ----------

waitForUser("Proceed with Analysis", 
    "You can close the graph.\n" +
    "Step 1: Delete the previous ROI.\n" +
    "Step 2: Draw new ROIs on the duplicated image, first cell length, then diagonal for each cell.\n" +
    "When ready, click 'OK' to proceed.");

// ------ Second Macro used for analysis and saving the output ----------


processedImage = outputPath + baseName + "processed_.tif";  // SAVES ONLY LATEST PICTURE 
saveAs("Tiff", processedImage);

// measure rois
roiManager("Show All");
roiCount = roiManager("count");

// prepare result table
run("Clear Results");

// loop through rois and process them based on the idex: odd for length
for(r = 0; r < roiCount; r++){
	roiManager("select", r); // select the roi
	if((r+1) % 2 == 1){
		// if roi is odd
		roiManager("select", r);
		roiManager("measure"); 
	}
}

// Get user input for genotype
genotype = getString("Enter sample label:", "genotype");


// Initialize count for the current genotype
count = 1;

for (i = 0; i < fileList.length; i++) {
    fileName = fileList[i];

    // Check if the filename matches this genotype and contains _length_
    if (endsWith(fileName, ".csv") && indexOf(fileName, "_length_") != -1 && startsWith(fileName, genotype + "_length_")) {
        parts = split(fileName, "_length_");
        if (parts.length > 1) {
            num = parseInt(replace(parts[1], ".csv", ""));
            if (!isNaN(num) && num >= count) {
                count = num + 1;
            }
        }
    }
}	

// Save cell length measurements
resultsPath = outputPath + genotype + "_length_" + count + ".csv";
saveAs("Results", resultsPath);
close("Results");

// for profiles, even numbers
run("Clear Results");

// Get the callibration
getPixelSize(unit, pixelWidth, pixelHeight, voxelDepth);

// For profiles, even numbers
profileIndex = 1; // Start numbering profiles from 1

for (r = 0; r < roiCount; r++) {
    if ((r + 1) % 2 == 0) {  // Select only even-numbered ROIs
        roiManager("select", r);
        profile = getProfile();

        // Clear previous results
        run("Clear Results");

        // Store profile values in the results table
        for (i = 0; i < profile.length; i++) {
            distance = i * pixelWidth; // Center of each pixel, in calibrated units
            setResult("Distance", i, distance);
            setResult("Value", i, profile[i]);
        }

        // Construct file name using genotype, count, and profileIndex
        profileResultsPath = outputPath + genotype + "_spectra_" + count + "_" + profileIndex + ".csv";

        // Save each profile in a separate CSV file
        saveAs("Results", profileResultsPath);
        close("Results");

        profileIndex++; // Increment profile number for the next even ROI
    }
}


// save ROIs 
roiSetPath = outputPath + baseName + "-rois.zip";
roiManager("Save", roiSetPath);

print("Processing complete! HURRA!!!! \n Your files are in: " + outputPath);
```
