## Additional file 1 for "The Tonoplast Topology Index - a new metric for describing vacuole organization": stage2_v1.ipynb

```
# %% [code]
# Load the necessaries libraries
import os
import pandas as pd
import numpy as np
import matplotlib.pyplot as plt
from scipy.signal import find_peaks
from scipy.ndimage import gaussian_filter1d

# %% [code]
######################## Function that import the csv from the current directory files as dataframes and store them as a DataFrame in a dictionary ######################## 

def import_csv_to_dataframes(directory):
    """ 
    Reads all CSV files from the current directory and store them as pandas DataFrames in a dictionary called dataframes.
    
    Args:
        directory (str): Path to the directory containing CSV files.
    
    Returns:
        dict: A dictionary where keys are filename (without extension) and values are pandas DataFrames.
    """
    
    files = os.listdir(directory)  # List all the files in the directory
    
    csv_files = [file for file in files if file.endswith('.csv')]  # Filter out CSV files
    
    dataframes = {} # Create empty dictionary to store the DataFrames
    
    for csv_files in csv_files:
        df_name = os.path.splitext(csv_files)[0] # Extract filename without extansion
        dataframe = pd.read_csv(os.path.join(directory, csv_files)) #Read CSV files
        dataframes[df_name] = dataframe # Store Dataframes in dictionary
    return dataframes
           

######################## Function to detect and plot peaks from the DataFrames containing _spectra_ ########################
def plot_profile(dataframes, background=0, med_ratio=1.4, min_ratio=1.4, distance=0.1, prominence=0.1, sigma=0.01):
    """
    Identifies peaks in spectral data and visualise the detected peaks.
    
    Args:
        dataframes (dict): dictionary of DataFrames containing spectral data.
        background (flaot): minimum background threshold for peak detection.
        med_ratio (float): median-based filtering ratio.
        min_ratio (float): minimum peak value filtering ratio.
        distance (float): minimum distance between peaks.
        prominence (float): minimum prominence of peaks.
        sigma (float): smoothing factor for Gaussian filter
    
    Returns:
        peaks_dict: dictionary containing peak information (indices, distance, and values) for each DataFrame.
    """
    peaks_dict = {}
    
    for key, df in dataframes.items():
        if 'spectra' in key:
            # Extract distance and intensity values
            distance_list = df['Distance'].tolist() 
            value = df['Value'].tolist()

            # Use the median intensity to define a threshold for valid peaks
            median_value = np.median(value)
            threshold = max(background, med_ratio * median_value)
            
            # Apply Gaussian smoothing to reduce noise and make peak detection more robust
            smoothed_value = gaussian_filter1d(value, sigma=sigma)
            
            # Identify peaks using scipy's find_peak function
            peaks, properties = find_peaks(smoothed_value, height=threshold, distance=distance, prominence=prominence)

            # Further filter peaks to exclude weak signal
            filtered_indices = [i for i in peaks if value[i] > med_ratio * median_value]
            filtered_indices = [i for i in filtered_indices if value[i] > min_ratio * min(value)]
            filtered_indices = [i for i in filtered_indices if value[i] > background]

            # Store peak indices, distance, and values in a dictionary
            peaks_dict[key] = {
                'peaks_indices': filtered_indices,
                'peaks_distances': np.array(distance_list)[filtered_indices].tolist(),
                'peak_value': np.array(value)[filtered_indices].tolist()
            }
            
            # Plot original and smothed signal with detected peak
            plt.figure(figsize=(10, 6))
            plt.plot(distance_list, value, label='Original Value', alpha=0.5)  
            plt.plot(distance_list, smoothed_value, label='Smoothed Value', linewidth=2)  # Smoothed signal

            
            # Scatter plot of the peaks with red crosses
            plt.scatter(np.array(distance_list)[filtered_indices], np.array(value)[filtered_indices], color='red', marker='x', label='Peaks')
            plt.title(f"{key} - Min: {min(value):.3f}, Median: {median_value:.3f}, Max: {max(value):.3f}, Peaks: {len(filtered_indices)}")
            plt.xlabel('Distance')
            plt.ylabel('Value')
            plt.legend()
            plt.grid(True)
            plt.show()

    return peaks_dict


######################## Function to generate the final output DataFrame from extracted peak information ########################
def populate_df_from_dict(final_df, dataframes, peaks_dict):
    """ 
    Populate the final output table 
    
    Args:
        final_df (pd.Dataframe): existing DataFrame created in the beginning to apend results to.
        dataframes (dict): dictionary of the imported csv files.
        peaks_dict (dict): dictionary containing peak detection results
    
    Returns:
    pd.DataFrame: Updated final_df with the information regarding the cells
    """
    rows = []
    for key in dataframes:
        if 'spectra' in key:
            parts = key.split('_')
            if len(parts) >= 4:
                genotype, spectra, sample, cell = parts[0], parts[1], parts[2], parts[3]
                
                # Look in the peak_dict for the correct key so I can add the peak count in the correct row
                peak_count = len(peaks_dict[key]['peaks_indices']) if key in peaks_dict else None
                
                # Calculate the Transect length as the last Distance value
                transect = None  
                if not dataframes[key].empty and 'Distance' in dataframes[key].columns:
                    transect = dataframes[key]['Distance'].iloc[-1]  # Take last value instead of max()
                    
                # Add the cell length value
                length_key = f"{genotype}_length_{sample}"
                length = None  
                if length_key in dataframes and not dataframes[length_key].empty:
                    cell_index = int(cell) - 1  # Convert cell to int
                    if cell_index < len(dataframes[length_key]):
                        length = dataframes[length_key].iloc[cell_index]['Length']
                    
                # Populate the empty df with new rows
                new_row = {
                    'Sample': genotype,
                    'Image': sample,
                    'Cell': cell,
                    'Length': length,
                    'TTI': peak_count / transect if transect else None,  # Avoid division by None
                    'Transect Length': transect,
                    'Peaks Count': peak_count
                }
                rows.append(new_row)
    
    new_df = pd.DataFrame(rows, columns=final_df.columns)
    result_df = pd.concat([final_df, new_df], ignore_index=True)
    
    # Sort the columns in order to have them from A to Z
    result_df = result_df.sort_values(by=['Sample', 'Image', 'Cell'], ascending=[True, True, True]).reset_index(drop=True)
    
    # Transform the peak columns to numeric
    result_df['Peaks Count'] = pd.to_numeric(result_df['Peaks Count'], errors='coerce')
    
    return result_df


# %% [code]
# Initialise the empty dataframe that will be used to store the final data
columns = ['Sample', 'Image', 'Cell', 'Length', 'TTI', 'Transect Length', 'Peaks Count']

final_df = pd.DataFrame(columns = columns)

# %% [code]
# Importing files
dataframes = import_csv_to_dataframes('.')
# List the keys of the dictionary
dataframe_keys = list(dataframes.keys())
print(dataframe_keys)

# %% [code]
# Variables for plot_profile function
background = 0
med_ratio = 0.9
min_ratio = 1.4
distance = 1
prominence = 0.1 # lower if valid peaks are missing
sigma = .01

# %% [code]
# find the peaks
peaks_dict = plot_profile(dataframes, background=background, med_ratio=med_ratio, min_ratio=min_ratio, distance=distance, prominence=prominence, sigma=sigma)

# %% [code]
# Create the final Dataframe

final_df = populate_df_from_dict(final_df, dataframes, peaks_dict)

# %% [code]
# Check if the data are randomised or not
plt.figure(figsize=(10,6))
plt.scatter(final_df['Length'], final_df['Transect Length'], c = 'blue')

# Add labels and titles
plt.xlabel('Cell length')
plt.ylabel('Transect length')
plt.title('Scatter plot of Transect Lenght vs Cell Length')

plt.grid(True)
plt.show()

# %% [code]
# Saving the output as excel
final_df.to_excel("output.xlsx")


# %% [code]
print(final_df)
```
