## Additional file 1 for "The Tonoplast Topology Index - a new metric for describing vacuole organization": TTI_manual_v1.pdf

### TTI – Tonoplast Topology Index

#### *Version 1.0 – user manual*

**Package contents:**

- This document
- ImageJ macro: `Stage1_v1.ijm`
- Jupyter Notebook file: `Stage2_v1.ipynb`

**Required:**

- A PC running a recent version of ***Fiji*** (http://fiji.sc/Fiji), ***Jupyter notebook*** (typically installed as a part of the ***Anaconda*** package, see Appendix 1) and a spreadsheet program such as ***Microsoft Excel*** or ***Libre Office Calc***.
  - *Performance under different OS or on different system configurations may vary but the program has been successfully used on a PC with Intel Core 5 @ 3.1 GHz and 16 GB RAM running the Fiji distribution of ImageJ 1.54p, Anaconda 2.5.1 and MS Excel under a 64bit version of Win10 Enterprise. The instructions described here refer to the Windows environment, but the program also works on Linux-based OS and was successfully tested under Ubuntu 24.04 and on an Apple Mac system.*
- Confocal microscopy images (preferably z-stacks) – either original files produced by the microscope´s software or standard `*.ome.tiff` files (8-bit images are OK). We are typically measuring atrichoblasts from isodiametric stage until the first cells with a large contiguous vacuole, which in our setup are about the length of ~1/2 screen.
  - *The workflow was not tested with multi-channel images, but it should be possible.*
  - *Images that do not carry correct and consistent scale information in their metadata can also be analysed if you define the scale manually in* ***Fiji*** *(e.g. by drawing a line of known size on a scale slide image and setting the scale as global).*

**Conventions in this manual:**

Instructions in plain text, file/folder names and variable/parameter names in `Courier`, program and menu names in ***bold italics***, commands in **bold**, program output quotes in **`Courier bold`**, notes, remarks and comments in *italics*.

**Before you begin:**

- Prepare a folder structure containing your image data. It is recommended to make a separate folder for each genotype or treatment (e.g., `WT` and `mutant`).
- Copy `Stage1_v1.ijm` into a location on your local drive where you can easily find it.

**Stage 1**

1. Open ***Fiji***.
2. Set the parameters to be measured: in the ***Fiji*** menu, go to ***Analyze*** – ***Set Measurements***, preferably uncheck everything and click **OK**.
   - *This needs to be done only once at the beginning or after doing something else in Fiji.*
   - *Leaving something selected should not affect the results, but bigger files containing unnecessary additional data and will be produced; nevertheless, they still should be suitable for Stage 2 processing.*

CC-BY-4.0 Helena Kočová, George Caldarescu and Fatima Cvrčková, 2025

3. Open the ***ROI manager*** (**Analyze – Tools – ROI Manager**). Check **Show all** and **Labels**.
4. Drag `Stage1_v1.ijm` onto the ***Fiji*** menu bar. This opens the interface of the macro.
5. Select the first file to be analyzed and open it by dragging onto the ***Fiji*** menu bar.
6. In the ***Bio-Formats import Options*** dialog box, select **View stack with Standard ImageJ**, **Stack order Default**, **Use virtual stack**, **Split channels**. Make sure no other boxes are checked and click **OK**. Your stack will open.
   - *If working with a multichannel image, close the unwanted channels at this step.*
7. Select an optical section in which the vacuole membranes are clearly visible in several cells but the optical section does not cross the nucleus, which is typically visible as a roughly circular void (in small cells with fragmented vacuole in the meristematic zone this may be difficult to distinguish). These cells will be measured.
   - *You can later go back and select another slice for measuring more cells; however, this will be handled as a separate sample and you will have to keep record separately to make sure you know which cells come from the same root.*
8. Draw a linear transect across a cell for checking plot quality. **Add** it to ***ROI Manager***.
9. **Run** the macro and check whether the resulting graph shows peaks corresponding to visible membrane crossings and none other. The default value of smoothing cycles is working fine with our SDCM images.
   - If you are not happy with the graph, press **Cancel**, close the graph and the duplicated single slice image that was generated (without saving). Run the macro again and experiment with the `Smooth` parameter until you get a satisfactory graph (you might need to do this step several times).
10. If you are happy with the graph, follow the instructions in the macro´s dialog window up to pressing **Proceed** and receiving the final message informing you about output file location.
    - You will have the opportunity to input sample name at this stage. In the following part of this document, we will assume that it is named `sample`, which is the default. Avoid using _ (underline) in the name, as it could break Stage 2 analysis.
11. You will find a new subfolder `Results` in your images folder, containing one summary file named `sample_length_x.csv` where `x` is the sample number (e.g. 1, 2, 3, etc) and multiple `sample_spectra_x_y.csv` files, where `x` is the same number as in `sample_length_x.csv` and `y` is the number of the analyzed cell, which corresponds with the row numbers in the `sample_length_x.csv` file. These files will be used for Stage 2.
    - *There will also be a zipped file with ROIs and an overlay image that you might find useful when troubleshooting. If you intend to reanalyze the same image stack, for example when going back to step 7 for another level of focus, rename these files, because they will get overwritten when you run the macro again (their filenames are automatically derived from the source image file name).*
12. To analyze next image, close everything except ***Fiji***, ***ROI Manager*** and the macro, delete all ROIs, open another image file and proceed to step 3. Repeat until you have processed all your files and then exit ***Fiji***.

**<u>Before Stage 2:</u>**

- Copy one instance of `Stage2_v1.ipynb` into each of the folders that contain your `*.csv` files from Stage 1.

  - If you have altered names of these files, make sure that thry follow this naming convention: `sample_length_x.csv` and `sample_spectra_x_y.csv` where `sample` is your sample name assigned in Stage 1 step 10, `x` is the plant (replica) and `y` is the cell label; `x` and `y` must be numbers (integers).
  - *CAUTION: the `*.ipynb` file will get altered in the process, keep a backup copy elsewhere.*

__Stage 2__

- Open `Stage2_v1.ipynb` in the folder containing your data from Stage 1 in ***Jupyter notebook*** (see Appendix 1 for instructions how to do it).
- Execute all blocks of code sequentially, examining output of each block for completeness and absence of error messages.
  - Make sure to manually examine the profile plots output to ascertain whether any false positive peaks have been detected (marked by a red x), or if there are false negatives, i.e. missed clearly visible peaks. Should this occur, it is necessary to adjust the parameters in the immediately preceding cell (marked **Variables for plot_profile function**) accordingly and re-run that cell before continuing. You may not be able to eliminate misdetection completely, but an occasional extra peak or missing peak in a small fraction of profiles will not affect the results substantially. Relevant parameters (with default values in brackets) are:
    - `background`: minimum intensity value to consider as peak (0)
    - `med_ratio`: a multiplier of the median value to set threshold detection (0.9)
    - `min_ratio`: a multiplier of the minimum value to filter peaks (1.4)
    - `distance:` minimum distance among separate peaks (1)
    - `prominence`: controls peak prominence, lower the value if missing peaks (0.1)
    - `sigma`: Gaussian smoothing; increase the value for noisy data possibly up to 1 (0.01)
  - At the penultimate block before saving the output, you will receive a scatter plot of transect length vs cell length. This serves as a data consistency check – the points should be located at, or close to, the diagonal. If you see any noticeable outliers, or for points that are still on diagonal but very far from others (indicating a possibly miscalibrated image file), check for errors and inconsistencies in your input data.
  - Any plots generated in the course of the analysis can be copied from the on-screen output as `*.png` if needed.
- At the end, results will be written into an Excel spreadsheet called `output.xlsx` in the same directory, which can be used for further statistical analysis.

CC-BY-4.0 Helena Kočová, George Caldarescu and Fatima Cvrčková, 2025

#### Appendix 1: *Jupyter notebook* basics for newbies

##### Setting up and starting *Jupyter notebook*

- ***Jupyter notebook*** is a part of the ***Anaconda*** package (https://www.anaconda.com/download).
    - If ***jupyter notebook*** is not installed, open ***anaconda prompt*** and type **conda install jupyter**. If asked for confirmation, type **y** and wait for installation to finish. You can also use the **conda install <package name>** command to install any missing package. Alternatively, you can use **pip install <package name>** in a python interface (such as ***jupyter notebook***). By default, ***python 3*** is used; if you have multiple python installations including ***python2***, use **pip3**.
    - The libraries required for running ***TTI*** should be included in the ***Anaconda*** installation, namely: ***os*** for file handling, ***pandas*** for data manipulation, ***numpy*** for numerical computations, ***matplotlib.pyplot*** for visualisation, ***scipy.signal*** for peak detection, ***scipy.ndimage*** for smoothing profiles.
- There are several possibilities to start ***Jupyter notebook***. The simplest are:
    - Open the ***Anaconda navigator*** app and open ***Powershell prompt*** from there. Navigate to the drive containing your `*.ipynb` file using shell commands, type **jupyter notebook** and press enter.
    - Open the ***Anaconda prompt*** from the Windows start menu. Navigate to the drive containing your `*.ipynb` file using shell commands, type **jupyter notebook** and press enter.

  A command line will appear followed by a **Jupyter notebook** tab in your default browser. ***Jupyter notebook*** runs locally and does not require internet access once installed. Do not close the command line (you can minimize it) as it would disconnect the ***Jupyter notebook***.
    - If you closed it accidentally and have unsaved data in your browser, you can renew it by starting another **Jupyter notebook** with a fresh command line.
- Navigate through the file system in the **Jupyter notebook** browser tab (single clicks to change directory), find your `*.ipynb` file and run it by single-clicking.

##### Working in *Jupyter notebook*

- ***Jupyter notebook*** is divided into multiple blocks (cells) of code, which you can run separately (unlike in other tools where you usually have to run whole code at once) – i.e., you are having a dialogue with the program, where you get an output of each step upon request.
- To run a block, locate cursor at its beginning and click the **Run** icon.
- During one session, it remembers all previous blocks of code you ran (unless overwritten), even if you deleted some code blocks in the meantime. However, this history is lost after restart (code blocks remain if you save the file).
- Some code blocks may take longer to process – wait until there is a number (for example **`In [1]`**) before running the next one, ***** means the block is still running. You may also notice the notebook icon in the browser changing into hourglass with longer code and the computer might become slow.
